# Heme dependent transcriptional regulation of the *Pseudomonas aeruginosa* tandem sRNA *prrF1,F2* locus by the cytoplasmic heme binding protein PhuS

**DOI:** 10.1101/2020.11.18.388926

**Authors:** Tyree Wilson, Susana Mouriño, Angela Wilks

**Affiliations:** Department of Pharmaceutical Sciences, School of Pharmacy, University of Maryland, Baltimore, MD 21201.

## Abstract

*Pseudomonas aeruginosa* is an opportunistic pathogen requiring iron for its survival and virulence. *P. aeruginosa* can acquire iron from heme via the heme assimilation system (Has) and *Pseudomonas* heme uptake (Phu) systems. The Has and Phu systems have non-redundant roles in heme sensing and transport, respectively. However, despite their respective roles heme taken up by either the Has or Phu system is regulated at the metabolic level by the cytoplasmic heme binding protein PhuS, which controls heme flux through the iron-regulated heme oxygenase HemO. Herein, through a combination of CHIP-PCR, EMSA and fluorescence anisotropy we show PhuS binds upstream of the tandem iron-responsive sRNAs *prrF1,F2.* Furthermore, qPCR analysis of the PAO1 WT and Δ*phuS* allelic strain shows loss of PhuS abrogates the heme dependent regulation of PrrH. Taken together our data shows PhuS, in addition to its role in regulating extracellular heme metabolism also functions as a transcriptional regulator of the heme-dependent sRNA, PrrH. This dual function of PhuS is central to integrating extracellular heme utilization into the PrrF/PrrH sRNA regulatory network critical for *P. aeruginosa* adaptation and virulence within the host.

## INTRODUCTION

Bacterial pathogens must acquire iron from their host for survival and virulence where due to its reactivity, it is tightly regulated and sequestered in iron-binding proteins such as transferrin and ferritin or in heme and iron-sulfur cluster containing proteins. Iron is further limited within the host during infection by the hosts’ innate immune response that includes the secretion of high affinity iron binding proteins such as lipocalin 2, and hepcidin dependent downregulation of plasma iron levels (1). To circumvent this nutritional immunity invading pathogens possess several acquisition strategies to acquire iron, and many encode systems for the utilization of heme (2–5). The gram-negative opportunistic pathogen *Pseudomonas aeruginosa* encodes two heme uptake systems; the heme assimilation system (Has) and the *Pseudomonas* heme uptake (Phu) system (6). The Has and Phu systems were shown to have non-redundant roles in heme sensing and transport, respectively (7). The Has system encodes an extracytoplasmic function (ECF) σ/anti-σ factor system, HasIS (6). ECF σ factors are a sub-family of alternative σ_70_ factors that allow for transcriptional amplification of genes involved in extracellular stress-response functions (8,9). The secreted hemophore HasAp on interaction with the OM receptor HasR triggers activation of the extracytoplasmic function (ECF) σ/anti-σ factor system HasIS. However, heme transported by either the HasR or PhuR outer membrane receptors is translocated to the cytoplasm by the *phu* encoded ABC transporter PhuUV, and its cognate periplasmic heme binding protein PhuT. Previous studies have shown the cytoplasmic heme binding protein PhuS regulates the flux of heme into the cell through a specific interaction with the iron regulated heme oxygenase, HemO (10,11). HemO oxidatively cleaves heme to release iron, CO and biliverdin (BV) IXβ and IXδ (12). Interestingly, the HemO metabolite BVIXβ is a post-transcriptional regulator of HasAp protein levels (13). Thus, the PhuS-HemO couple regulates both the flux of heme into the cell, and the extracellular heme signal through the heme metabolites BVIXβ and IXδ.

The complexity of *P. aeruginosa* iron and heme homeostasis is further exemplified by the tandem arrangement of the PrrF1 and PrrF2 sRNAs found directly downstream of the *phu* operon (Fig 1A) (14). The PrrF sRNAs are highly homologous to one another and contribute to iron homeostasis by causing mRNA degradation of non-essential iron-containing proteins (14–16). The PrrF sRNAs play a role in numerous other processes including twitching motility, quorum sensing molecule production, and biofilm formation. Furthermore, this tandem arrangement allows for the expression of an overlapping noncoding RNA, PrrH, whose expression is heme-dependent (17). The duplication of the *prrF* genes and the presence of *phuS* are genetically linked, and found in pathogenic *P. aeruginosa* but not in other Pseudomonads (17). Interestingly, PrrH is detected in infected murine lungs as well as sputum from cystic fibrosis (CF) patients suggesting a role for this sRNA during infection (18). In addition deletion of *phuS* or the *prrF1,F2* locus gave similar iron dysregulation transcriptomic profiles (15,19). Previous studies have shown the *S. dysenteriae* ShuS, a homolog of PhuS, has DNA binding properties (20). Based on these previous studies, we hypothesized that PhuS may also possess DNA binding properties providing a functional link between PhuS and the *prrF1,F2* locus. To test this hypothesis we performed a series of *in vivo* and *in vitro* experiments to determine the functional link between PhuS and *prrF* locus. Herein, through ChIP-PCR, EMSA and fluorescence anisotropy (FA) we show that apo-PhuS binds with high-affinity to the promoter of *prrF1* but not that of *prrF2*. Furthermore, comparison of the relative expression of PrrF and PrrH in the PAO1 WT and Δ*phuS* allelic strains by qPCR shows a loss in the heme dependent regulation of PrrH in the absence of PhuS. We propose PhuS has a dual function integrating extracellular heme metabolism into the iron-homeostasis networks through transcriptional modulation of the PrrF/PrrH sRNAs.

**Figure 1.**
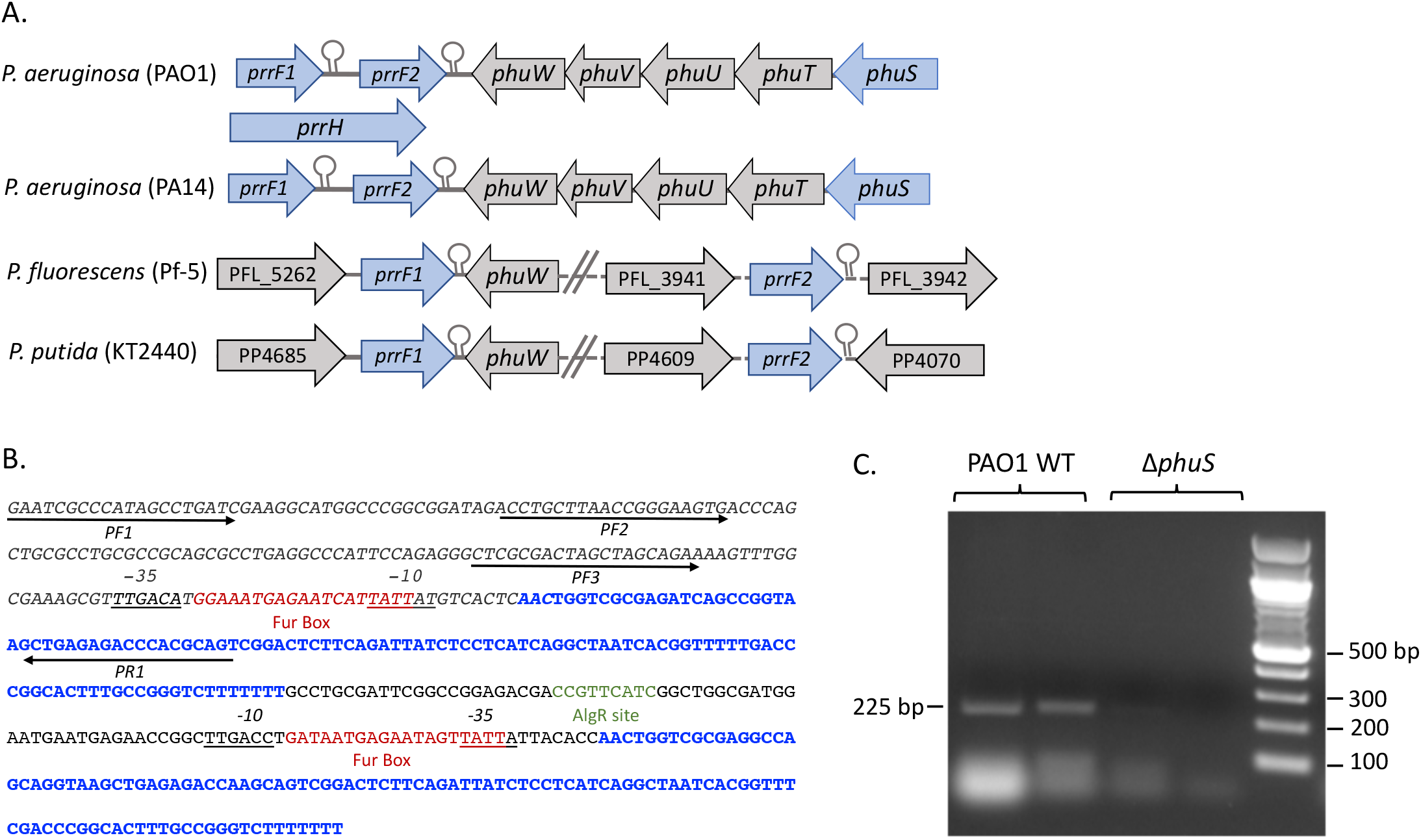
ChIP-PCR analysis of PhuS binding to the *prrF1,F2* promoter. *A.* Genetic organization of the *prrF1,F2* locus in pathogenic and non-pathogenic *Pseudomonas* strains. *B.* Sequence of the *prrF1,F2* locus. PrrF1 and PrrF2 are shown in blue. The Fur boxes upstream of *prrF1* and *prrF2* are shown in burnt red, the AlgR site in green and the −35 and −10 sites are underlined. Primers used in ChiP-PCR pull downs are indicated by the black arrows. The italicized sequence represents the fragment obtained following ChIP-PCR and DNAse I treatment. *C*. PCR fragments (225 bp; utilizing primers PF1 and PR1) isolated from PAO1 WT cells or Δ*phuS* control strain following crosslinking and pull-down with anti-PhuS. Lane 1. PAO1 WT in iron-deplete media; 2. PAO1WT supplemented with 1 μM heme; 3. Δ*phuS* iron-deplete in media; 4. Δ*phuS* supplemented with 1 μM heme; 5. DNA markers as shown. Bands were visualized on 1% agarose with ethidium bromide staining.

## RESULTS

### Isolation of a PhuS-prrF1 complex by ChIP-PCR

The potential for PhuS to bind to the *prrF1* promoter was analyzed by subjecting *P. aeruginosa* (PAO1) WT or the Δ*phuS* deletion strain to chromatin immunoprecipitation (ChIP) followed by PCR amplification. Employing primer pairs that specifically amplify the promoter region of the *prrF1* promoter (Table S1). Following DNAse digestion, reversal of the formaldehyde cross-linking and sequencing we determined a ~230 bp fragment upstream of the *prrF1/prrH* promoter that includes part of the Fur box sequence (Fig 1B). We repeated the pull-downs with purified genomic DNA (100-500 bp sheared fragments) and addition of purified His-tagged PhuS (PhuS-His6) to PAO1 WT. Ni-NTA pull down, DNAse treatment and PCR amplification with the primer pairs shown in Fig 1A identified bands of ~230, 180 and 120 bp which following sequencing confirmed binding to the *prrF1* promoter (Fig S1). Due to the high sequence identity (>95%) between PrrF1 and PrrF2 we were unable to design primers to specifically probe PhuS binding to the *prrF2* promoter. PhuS binding to the *prrF2* promoter was analyzed by FA (see following section).

### Apo-PhuS and Fur have overlapping binding sites within the prrF1 promoter

Interestingly, the DNA fragment isolated by ChIP-PCR included the *prrF1* Fur-box (Fig 1B). Utilizing FA we analyzed Fur and PhuS binding to a 5’fluoroscein (5’-FAM) labeled 30 bp oligonucleotide encoding the Fur box alone (*prrF1*-30) (Table S2 and Fig 2A). The change in anisotropy on addition of Fur when fit to a one to one binding site model gave a binding constant (KD) of 50 ± 10 nM (Fig 2A). In contrast addition of apo-PhuS to *prrF1*-30 showed a much smaller change in anisotropy with a K_D_ >2 μM (Fig 2B). However, titration of a 5’-FAM labeled oligonucleotide that includes sequence upstream of the Fur box (*prrF1*-50) significantly enhanced PhuS binding (Table S2 and Fig 2B). The change in anisotropy when fit to a one to one binding site model gave a K_D_ of 64 ± 10 nM (Fig 2B). In contrast a 5’-FAM labeled oligonucleotide encompassing the upstream sequence but lacking the Fur box showed no change in anisotropy (Table S2 and Fig S2). Therefore, the optimal binding of PhuS to the *prrF1* promoter requires sequence upstream of and including the Fur box. To confirm PhuS specificity for the *prrF1* promoter over that of *prrF2* we performed FA on 5-FAM labeled oligonucleotides designed within the *prrF1,prrF2* intergenic region (Table S2). Incremental addition of apo-PhuS to 5’-FAM labeled oligonucleotides *prrF*2-50 (Fur) including the Fur box or the upstream *prrF*2-50 (AlgR) which includes the AlgR site (Fig 1A and Table S2) showed no change in anisotropy (Fig S2), confirming PhuS specificity for the *prrF1* promoter.

**Figure 2.**
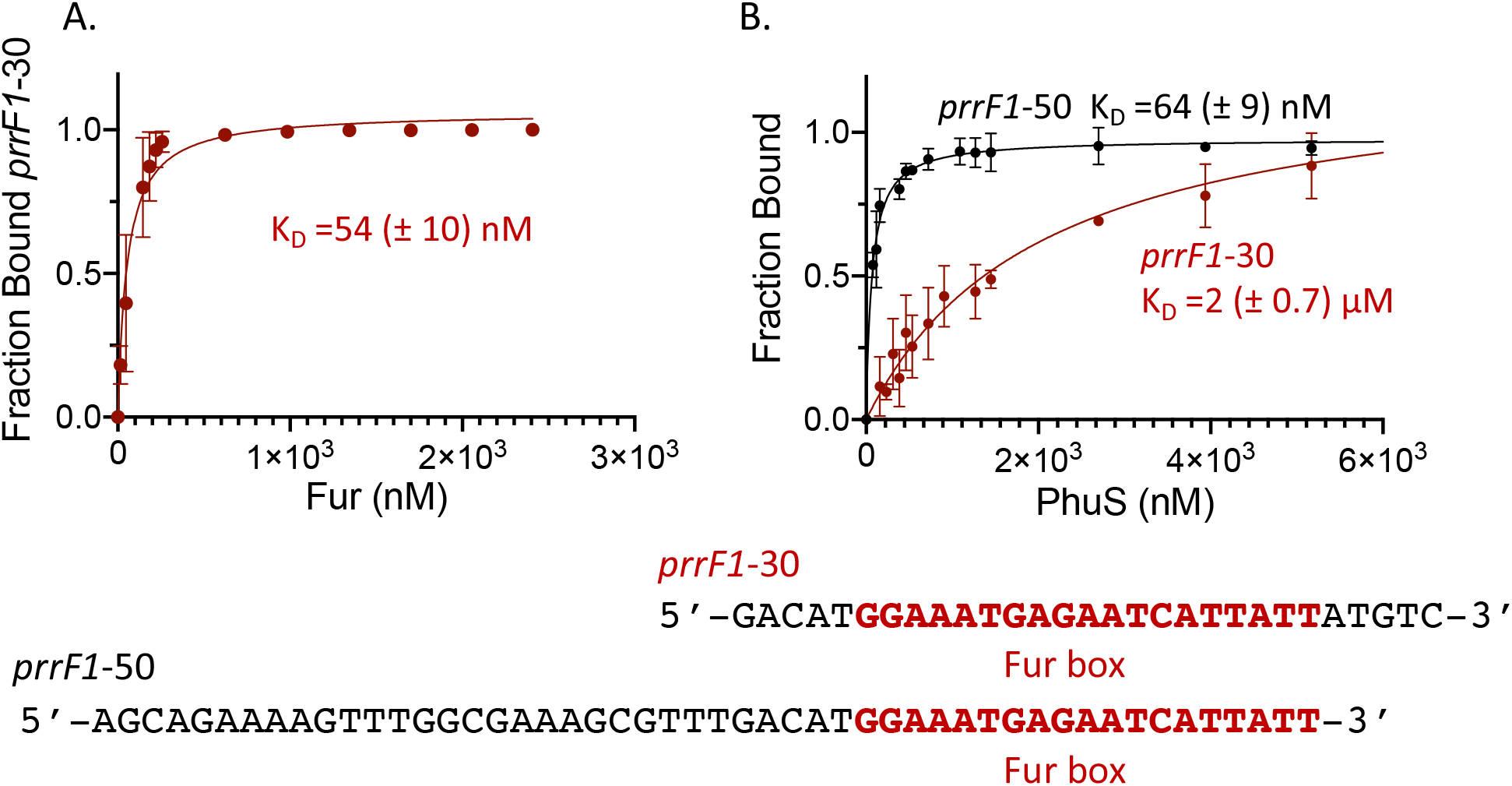
Fluorescence anisotropy of Fur and PhuS binding. *A.* Mn-Fur binding to the 5’-FAM labeled *prrF1*-30. *B*. apo-PhuS binding to the 5’-FAM labeled *prrF1*-50 and *prrF1*-30 color coded as shown. Experiments were performed in triplicate as described in Experimental Procedures. The data was fit by converting the anisotropy, *r*, to fraction bound and plotted against protein concentration using a one-site binding model. The error is shown as the standard error of the mean (SEM).

### Apo-PhuS but not holo-PhuS binds to the prrF1 promoter

Following characterization of the PhuS binding region we next sought to determine if heme and DNA binding were mutually exclusive. In contrast to apo-PhuS the addition of holo-PhuS to the 5’-FAM labeled *prrF1*-50 oligonucleotide showed no change in anisotropy (Fig S2). The FA analysis was confirmed with EMSAs of apo- and holo-PhuS binding to a 5’biotinylated *prrF1*-50 oligonucleotide. Addition of increasing concentrations of apo-PhuS gave a lower mobility complex consistent with apo-PhuS binding to *prrF1*-50 (Fig 3A). In contrast addition of holo-PhuS showed no shift in biotinylated-*prrF1*-50 (Fig 3B). Taken together the data is consistent with heme and DNA binding being mutually exclusive functions of PhuS.

**Figure 3.**
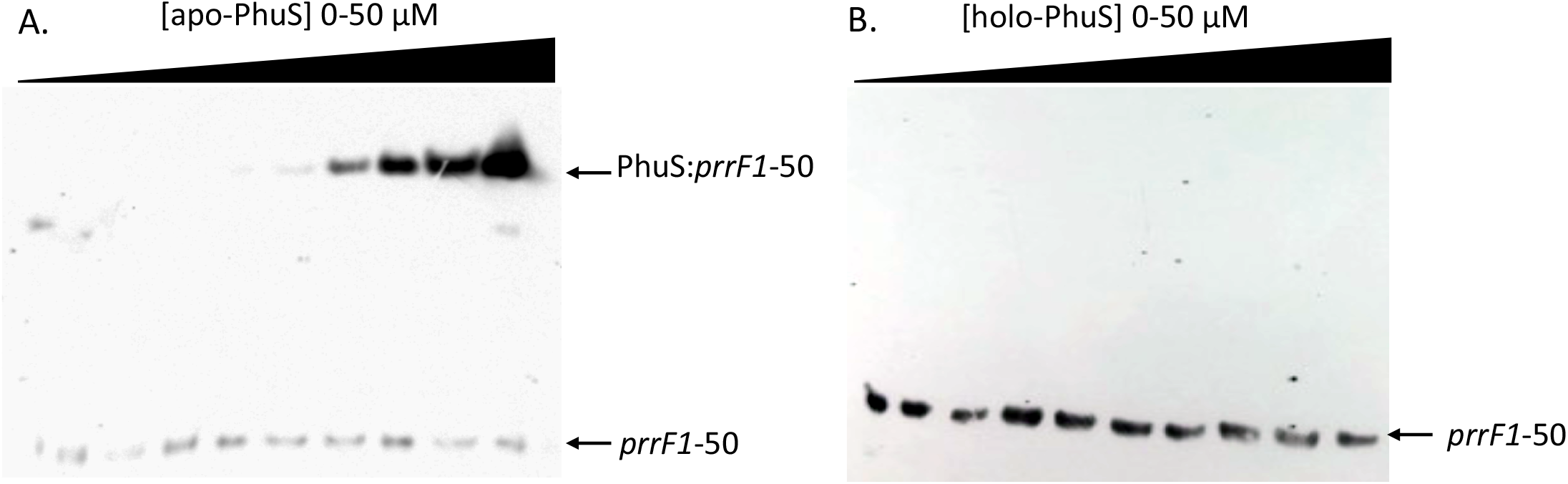
EMSA of apo- and holo-PhuS binding to *prrF1-50*. *A.* apo-PhuS binding to 5’-biotin labeled *prrF1*-50. *B.* holo-PhuS binding to 5’-biotin labeled *prrF1*-50. Experiments were performed as described in Experimental Procedures. All reactions contained a fixed concentration (30 pM) of labeled *prrF1*-50 and following incubation were run on 8% acrylamide gels and transferred to a nylon membrane and visualized by chemiluminescence.

### HemO modulation of the holo-PhuS to apo-PhuS equilibrium drives DNA binding

Our previous studies characterized PhuS as a titratable regulator of heme flux through HemO (11). Based on these studies we hypothesized heme flux through HemO may be coupled to PhuS regulation of the *prrF1,2* operon. We tested the ability of HemO to drive PhuS DNA binding by FA and EMSA. On titration of a fixed concentration of holo-PhuS and 5’-biotinylated *prrF1*-50 with increasing concentrations of apo-HemO we observe a lower mobility complex, consistent with apo-PhuS binding to *prrF*1-50 (Fig 4A). Similarly, on titration of a fixed concentration of holo-PhuS (1 μM) and 5’FAM-labeled *prrF1*-50 (10 nM) with HemO we observe an increase in anisotropy. The change in anisotropy when fit to a one to one binding site model shows a similar 1:1 saturation observed for the EMSA (Fig 4B).

**Figure 4.**
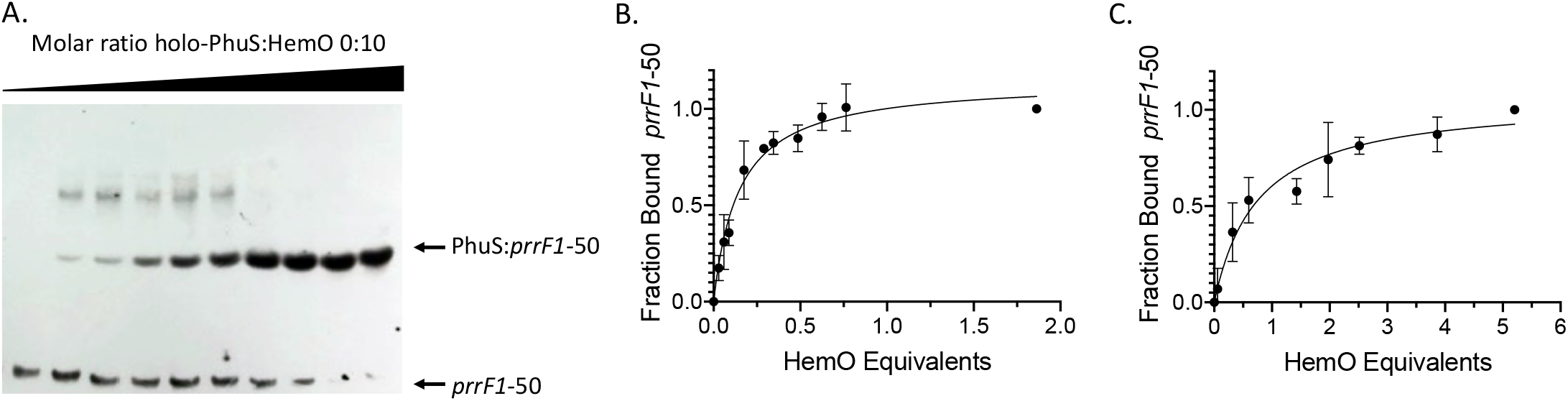
Holo-PhuS titration with apo-HemO drives DNA binding. *A.* EMSA of holo-PhuS titration with apo-HemO. Biotin labeled *prrF1*-50 (30 pM) and holo-PhuS (10 μM) was titrated with increasing concentrations of HemO (0-10 equivalents). Experiments were performed as in Figure 2. *B.* FA of holo-PhuS titration with *apo-HemO.* FA was performed with a fixed concentration of holo-PhuS (1 μM) and 5’-FAM labeled *prrF1*-30 (10 pM). The change in anisotropy was recorded as a function of apo-HemO molar equivalent until no further changes in anisotropy were recorded. *C*. As in B for holo-PhuS H212R. Experiments were performed in triplicate as described in the Experimental Procedures. The data was fit by converting the anisotropy, *r*, to fraction bound and plotted against HemO molar equivalents using a one-site binding model. The error is shown as the standard error of the mean (SEM).

We have previously shown that apo-PhuS undergoes a significant conformational rearrangement on heme binding (21), which likely accounts for the mutually exclusive roles in DNA binding and heme transfer. Furthermore, through site directed mutagenesis and spectroscopic studies we proposed a model where a conformational rearrangement on protein-protein interaction triggers a ligand switch within PhuS (from H209 to H212) prior to release to HemO. Furthermore, while the apo-PhuS H212R mutant rapidly binds heme the rate of heme transfer to HemO is inhibited (21). We sought to determine if the altered heme binding and transfer properties of this PhuS H212R mutant influenced binding to the *prrF1* promoter. We confirmed by FA that apo-PhuS H212R binds to 5’-FAM labeled *prrF1*-50 with a K_D_ of 90 ± 30 nM (Fig 5A). Although we saw a decrease in the binding affinity and fraction bound, EMSA analysis showed a shift consistent with complex formation (Fig 5B). Titration of a fixed concentration of holo-PhuS H212R and 5’FAM-labeled *prrF1*-50 (10 nM) with HemO drives apo-PhuS binding. However, consistent with the inhibition of heme transfer a significantly greater molar ratio of HemO to holo-PhuS H212R (5:1) is required to drive the reaction to completion when compared to holo-PhuS WT (Fig 4C).

**Figure 5.**
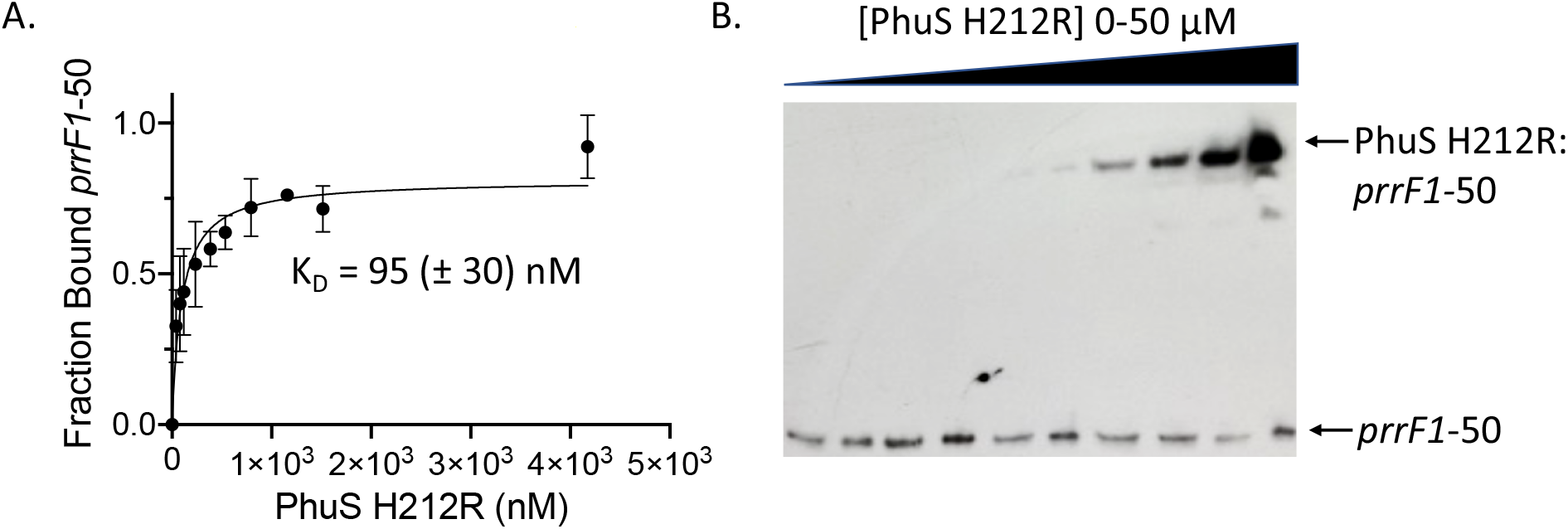
FA and EMSA of apo-PhuS H212R binding to *prrF1* **-50.** A. FA of apo-PhuS H212R binding to the 5’-FAM labeled *prrF1*-50 as in Fig 2. The data was fit by converting the anisotropy, *r*, to fraction bound and plotted against protein concentration using a one-site binding model. The error is shown as the standard error of the mean (SEM). *B.* apo-PhuS H212R binding to 5’-biotin labeled *prrF1*-50 as in Fig 3. All reactions contained a fixed concentration (30 pM) of labeled *prrF1*-50 and following incubation were run on 8% acrylamide gels, transferred to a nylon membrane and visualized by chemiluminescence.

### Heme flux through PhuS regulates PrrH but not PrrF1 levels in vivo

To assess the role of PhuS in transcriptional regulation of *prrF1* and/or *prrH* we performed qPCR analysis of PAO1 WT, a *phuS* knockout (Δ*phuS*) and the *phuSH212R* allelic strain in low iron and heme supplemented conditions. All of the strains utilized had similar growth rates in iron-deplete or heme supplemented conditions (Fig S2). Through a combination of isotopic ^13^C-heme uptake followed by LC-MS/MS and ICP-MS we have previously shown 1 μM heme supplemented cultures deplete the exogenous heme by 7-8 h, resulting in Fur repression (13,22,23). Therefore, we analyzed the relative PrrF and PrrH levels at 2 and 5 h where heme flux through PhuS is maximal and prior to Fur repression. The PrrF probe (Table S2) detects PrrF, PrrF2 and PrrH sRNAs due to the similarity and overlap in sequences. In contrast the PrrH probe comprising the unique intergenic sequence between *prrF1* and *prrF2* detects PrrH specifically (Table S2). Given the previously reported low abundance of PrrH compared to PrrF1 and PrrF2 (17), the contribution of PrrH to the relative RNA levels measured with the PrrF probe is negligible. In iron-deplete conditions we see a ~2-fold increase in PrrF at 5 h consistent with iron-deprivation (Fig 6A; left panel). However, in heme supplemented conditions at 2 h we observe an initial ~2-fold decrease in the relative PrrF levels. However, at 5 h the relative expression of PrrF on heme supplementation is identical to that in low iron conditions (Fig 6A; left panel). We have previously observed a similar heme dependent decrease in relative RNA levels at the early 2 h time point for Fur regulated genes within the *has* and *phu* operons (13,22). We attribute this decrease to an initial effect of the influx of heme or iron. In contrast to PrrF, PrrH levels show no increase over time in low iron (Fig 6A; right panel). However, in heme supplemented conditions we see a significant 3-4 fold increase in PrrH levels at 5h (Fig 6A; right panel). Furthermore, at the 2 h time point we do not observe the initial decrease in the relative expression of PrrH as seen for PrrF. Therefore, in contrast to PrrF which is under transcriptional regulation of Fur, PrrH is not iron-regulated but is subject to positive regulation by heme.

**Figure 6.**
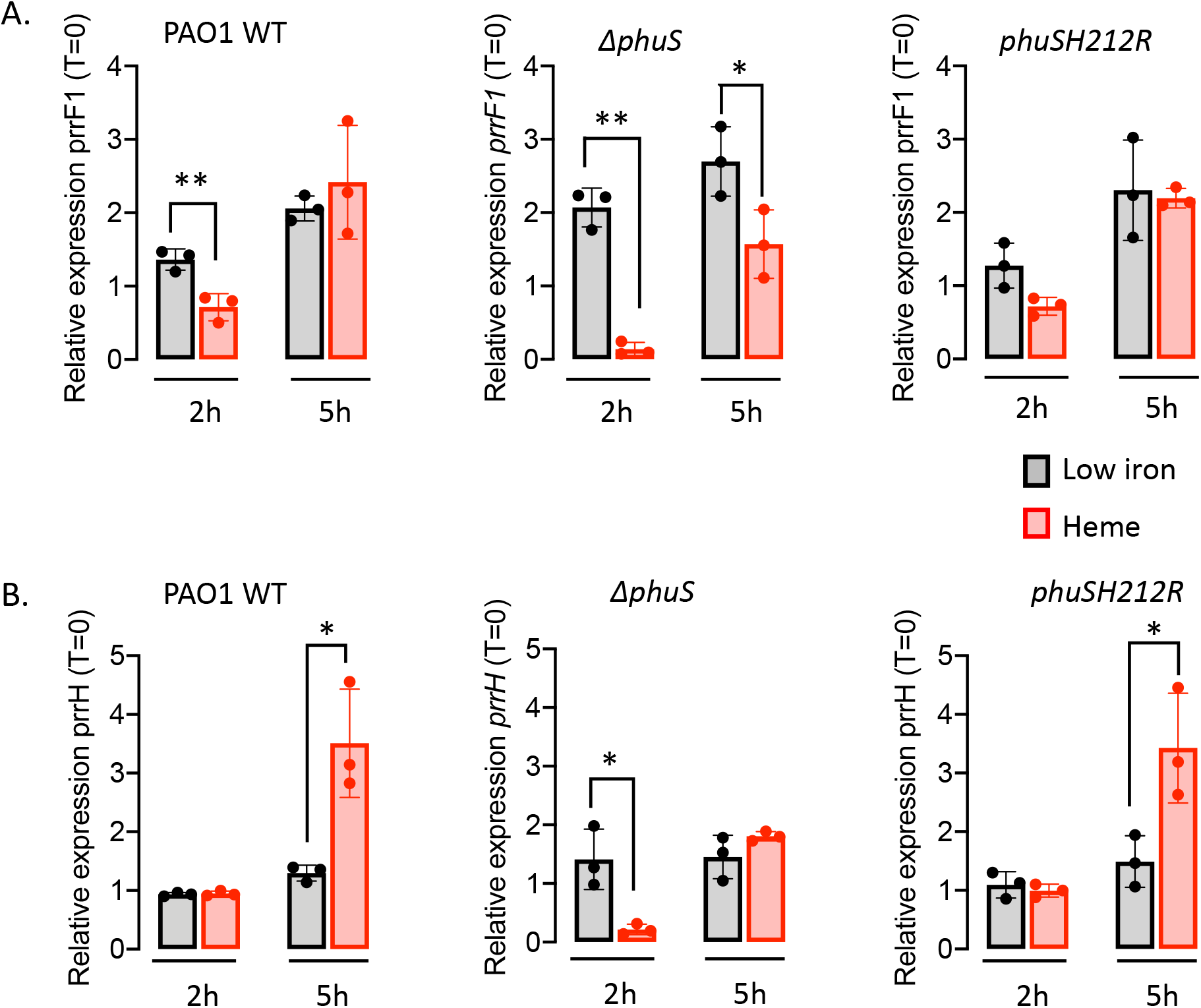
Relative PrrF1 and PrrH sRNA levels for PAO1 WT, *ΔphuS* and the *phuSH212R* allelic strain in iron-deplete or heme supplemented conditions. *A*. PrrF1 relative sRNA levels. *B*. PrrH relative sRNA levels. mRNA isolated at 0, 2, an 5 h following growth in either iron-deplete M9 or M9 supplemented with 1μM heme. mRNA values represent the mean from three biological experiments each performed in triplicate and normalized to 0 h. Grey shaded bars iron-deplete conditions; Red shaded bars heme supplemented conditions. *Error bars* represent the standard deviation from three independent experiments performed in triplicate. P values as determined by two-tailed Student *t* test comparing values upon heme supplementation to iron deplete conditions at the same time point, where *, *p* < 0.05.

To confirm the heme-dependent increase in PrrH expression is indeed mediated by PhuS, we performed qPCR analysis on the Δ*phuS* strain. In low iron we observe a ~2-fold increase in PrrF expression at the earlier 2 h time point in the Δ*phuS* strain (Fig 6A; middle panel). Interestingly, the decrease in relative expression of PrrF in heme supplemented conditions compared to low iron is significantly enhanced in the Δ*phuS* strain compared to PAO1 WT (Fig 6A; middle panel). On heme supplementation we observe a loss in the heme dependent regulation of PrrH in the Δ*phuS* strain (Fig 6B; middle panel). Deletion of PhuS not only leads to a loss in the heme-dependent increase PrrH expression, but also appears to increase the iron-effect over the *prrF1* promoter. Given the partial overlapping binding sites of PhuS and Fur, it is possible the absence of PhuS may allow for increased access to the Fur-box and enhanced repression of PrrF in the Δ*phuS* knockout. Although not statistically significant, in low iron the loss of PhuS appears to show slightly elevated levels of PrrF relative to PAO1 WT (Fig 6A). Taken together the data suggests that PhuS binding as a function of heme status regulates the relative expression of PrrF1 and PrrH through modulation of Fur binding.

Based on the increased molar ratio of HemO required to drive PhuS H212R binding to the *prrF1* promoter *in vitro*, we sought to determine if such changes in heme binding and transfer influence PrrH expression. The relative expression profiles of both PrrF1 and PrrH in the *phuSH212R* allelic strain were very similar to that of PAO1 WT in both low iron and heme (Fig 6). Taken together the decrease in binding affinity and shift in the molar ratio of HemO required to drive apo-PhuS DNA binding *in vitro* are not significant enough to disrupt the PhuS-HemO equilibrium *in vivo.* Future studies with variants disrupting either heme transfer or the PhuS-HemO protein-protein interaction will be undertaken to investigate the role of extracellular heme flux on PrrF and PrrH expression.

## DISCUSSION

Iron acquisition and homeostasis are critical for *P. aeruginosa* survival and pathogenesis. Bacterial iron homeostasis is maintained by either repressing the expression of iron-uptake systems in iron-replete conditions, or by decreasing the levels of iron-containing proteins in iron-limiting conditions. The latter function is most often mediated at the post-transcriptional level by iron-responsive sRNAs, that in many cases also regulate virulence traits (14,24–27). In *P. aeruginosa* the PrrF sRNAs play a role in numerous other processes including twitching motility, quorum sensing molecule biosynthesis, and biofilm formation (16,18). However, the regulatory mechanisms by which *P. aeruginosa* adapts to a particular iron source are not as well understood. For example, in chronic infection *P. aeruginosa* decreases its reliance on siderophores, while simultaneously increasing reliance on heme (28,29). This increased dependence on heme coincides with the upregulation of the Phu heme uptake system.

The heme uptake systems like their siderophore counterparts are globally regulated by the master regulator Fur but must also have additional levels of regulation that allow for a coordinated transcriptional response to heme. Some years ago it was reported the tandem arrangement of *prrF1* and *prrF2* allows for expression of a longer heme responsive sRNA PrrH, predicted to affect the expression of genes related to heme homeostasis (17). Given the genetic link between the *prrF1,F2* locus and *phuS* we hypothesized that heme flux through PhuS may play a role in integrating heme metabolism into the sRNA regulatory network. Herein, we show apo-PhuS specifically binds within the *prrF1* promoter and modulates the expression of PrrF and PrrH as function of extracellular heme flux.

The fact that optimal apo-PhuS binding includes the Fur box (Fig 2B), but has no affinity for the Fur box alone (Fig 2C), suggests the PhuS and Fur binding sites are not mutually exclusive but may be antagonistic. This is supported in part by the qPCR data where in iron-limiting conditions the absence of PhuS increases the relative levels of PrrF compared to PAO1 WT (Fig 6A). In contrast, in heme supplemented conditions the initial iron-dependent repression of PrrF is significantly increased, presumably a consequence of greater access of Fur to the Fur box in the absence of apo-PhuS (Fig 6A). The increase in Fur repression on loss PhuS also leads to a decrease in the relative expression of PrrH in heme supplemented conditions. These studies suggest apo-PhuS binding to the *prrF1* promoter as a function of Fur antagonism allows for a coordinated iron and heme transcriptional response. A previous study has shown the *prrF* operon requires extended upstream sequence for full promoter activity (18), a common feature of promoters that bind multiple transcription factors as higher order oligomers, and show promiscuous DNA shape dependent binding at sites distant from the transcriptional start site (30,31). The Fur proteins themselves are known to oligomerize in a metal-dependent manner and bind to promoters at multiple sites causing DNA looping (31–33). Similarly, the PhuS homolog ShuS was shown by atomic force microscopy (AFM) techniques to form oligomeric complexes condensing the DNA (20). The fact the PhuS protected fragment (Fig 1B) is ~200 bp is consistent with PhuS having similar nucleoid associated protein like properties that include oligomerization and promiscuous binding specificity. It is not clear at the present time how modulation of PhuS and Fur-binding to the *prrF1* promoter allows for re-modelling of the DNA structure or read through of the *prrF1* transcriptional terminator required for PrrH expression. Interestingly, upstream of the *prrF2* Fur-box is an AlgR binding site that has been shown to directly and indirectly regulate pyoverdine biosynthesis (34). The AlgR transcriptional regulator is part of the *algZR* two-component sensor system that regulates alginate as well as several virulence factors including Type IV pillus, rhamnolipid production, Rhl quorum sensing system, and biofilm formation (35–39). It is possible given the proximity of the *prrF1* and *prrF2* promoters that are separated by only 95 bps that short range DNA interactions driven by higher order multimers or overlapping interactions of the transcriptional regulators allows for differential expression of PrrF1, PrrF2 and/or PrrH. A precedent for such a mechanism has been characterized in *Helicobacter pylori* where oligomerization and DNA condensation by Fur, and its antagonism by the Ni-dependent NikR allows for integration of metal-homeostasis and acid acclimation (31). A more extensive analysis by DNAse I footprinting and expression analysis under different conditions will determine if these transcriptional regulators physically interact and coordinate transcriptional regulation via structural changes within the *prrF1* and *prrF2* promoters.

Differential regulation over the *prrF1* and *prrF2* promoters provides a mechanism by which the relative expression levels of PrrF1, PrrF2 and PrrH may ultimately determine the distinct target profiles of the sRNAs. This is especially true of PrrF1 and PrrF2 that have almost identical sequences. Interestingly, in PAO1 *algR* is co-transcribed with the *hemCD* genes providing a link to intracellular heme biosynthesis (40). Furthermore, the AlgR regulation of pyoverdine provides a link between heme biosynthesis and iron homeostasis (34). While specific targets of PrrH are not as well characterized the identification of potential PrrH targets such as *vreE* a regulator of virulence, the heme-d_1_ biosynthesis gene *nirL*, and the *phuS* gene itself, suggest PhuS dependent modulation of PrrH (similar to AlgR) may further allow for integration of iron and heme homeostasis with the virulence networks of *P. aeruginosa* (16,17). Indeed, the Δ*prrF1,F2* mutant is defective for both heme and iron homeostasis and is attenuated for virulence in an acute mouse lung infection model (18). Therefore, it is reasonable to suggest that the modulation of the iron-dependent PrrF/PrrH network by PhuS and AlgR plays a role in unifying intracellular iron and heme homeostasis as well as virulence traits required for infection. The ability to rapidly respond and adapt to heme as an iron source is likely to provide a competitive advantage in the host. As previously mentioned, in chronic infection *P. aeruginosa* adapts over time to utilize heme as an source iron via the Phu system, while decreasing its reliance on siderophore systems (28). The fact the tandem arrangement of the *prrF* genes and the presence of *phuS* are genetically linked, and found only in pathogenic *P. aeruginosa* highlights the significance of the iron and heme-dependent sRNAs in this adaptive response (17). Furthermore, the detection of both PrrF and PrrH in infected murine lungs as well as sputum from CF patients further signifies a role for these sRNAs during infection (18).

In summary, we have shown PhuS in addition to its role in regulating extracellular heme metabolism also functions as a transcriptional regulator of the heme-dependent sRNA, PrrH. This dual function of PhuS is central to integrating extracellular heme utilization into the PrrF/PrrH sRNA regulatory network critical for *P. aeruginosa* adaptation and virulence within the host. Based on these preliminary studies PhuS offers an advantage as a potential antimicrobial target; i) it is found only in pathogenic *P. aeruginosa* strains and ii) its dual function in pathways central to survival and pathogenesis in the host is potentially advantageous in slowing resistance development. A more complete understanding of the molecular mechanisms by which PhuS regulates a coordinated transcriptional response from the *prrF1* promoter will be critical in the development of novel strategies to target iron homeostasis and virulence.

## EXPERIMENTAL PROCEDURES

### Bacterial strains and growth conditions

Bacterial strains and plasmids used in this study are listed in Table S1 and oligonucleotide primers and probes in Table S2. All primers and probes used in this study were purchased from Integrated DNA Technology (IDT). *E. coli* strains were routinely grown in Luria Bertani (LB) broth (*American Bioanalytical*) or on LB agar plates. *P. aeruginosa* strains were freshly streaked and maintained on *P.* isolation agar (PIA) (*BD Biosciences*). All strains were stored at −80°C in LB with 20% glycerol. The iron levels in M9 medium (*Nalgene*) were determined by inductively coupled plasma-mass spectrometry to be less than 1 nM. For qPCR, singly isolated colonies from each *Pseudomonas* strain were picked, inoculated into 10 mL of LB broth, and grown overnight at 37 °C with shaking (210 rpm). The bacteria were then harvested and washed in 10 mL of M9 minimal medium. Following centrifugation, the bacterial pellet was resuspended in 10 mL of M9 medium and used to inoculate 50 mL of fresh M9 iron-deplete medium to a starting *A*600 of 0.04. Cultures were grown at 37 °C with shaking for 3 h before the addition of supplements (0 h) and incubated for a further 6 h. When required, antibiotics were used at the following final concentrations tetracycline (Tc) 10 and 150 μg mL^−1^ for *E. coli* and *P. aeruginosa*, respectively. When required, ampicillin (Amp) was used at a final concentration of 100 μg/mL.

### Construction of the Pseudomonas aeruginosa phuSH212R allelic strain

*phuSH212R* was obtained by allelic exchange as previously described (41), using the parental strain PAO1 Δ*phuS* (42). Briefly, a 2.9 kb *phuS* gene fragment including upstream and downstream sequence was PCR amplified from the chromosomal DNA of *P. aeruginosa PAO1* using primers pairs *Pst*I-5’PhuS-F and *Hind*III-3’PhuS-R. The amplified fragment was cloned into pUC18, resulting in pUC18-5’-PhuS-3’. The mutant allele *phuSH212R* was obtained following digestion of plasmid pET21*phuS*H212R (21) with *Nru*I and *Stu*I and subcloned into *Nru*I and *Stu*I digested pUC18-5’-PhuS-3’, replacing the wild type allele. The new construct pUC18-5’-*phuS*H212R-3’ was confirmed by sequencing (*Eurofins MWG Operon*). The insert including *phuS*H212R plus the 5′ and 3′ flanking regions was purified by *PstI*-*Hind*III digestion and ligated into the counter-selective suicide plasmid pEX18Tc (41). Finally, plasmid pEX18Tc-5’-phuSH212R-3’ was transferred into *P. aeruginosa* Δ*phuS* by conjugation. A double event of homologous recombination followed by selection on PIA plates containing 5% sucrose, resulted in chromosomal integration of *phuS*H212R, replacing the parental allele Δ*phuS.* PCR and sequencing analysis were used to verify the allelic exchange process.

### Expression and Purification of apo-PhuS, and PhuSH212R

Protein expression was performed as previously reported with slight modification(10,43). The PhuS or PhuS H212R mutant lysate was applied to a Sepharose-G column (*GE Life Sciences*) equilibrated with 20 mM Tris-HCl (pH 8.0) and washed with 5 column volumes of the same buffer. The column was further washed with 10 column volumes of 20 mM Tris (pH 8.0) containing 20 mM NaCl and the PhuS protein eluted with a linear gradient of 50-500 mM NaCl. Eluted fractions were analyzed by SDS-PAGE, and the peak fractions pooled and dialyzed against 4L of 20 mM Tris (pH 8.0) containing100 mM NaCl. The protein was concentrated in a Pierce™ Protein Concentrator (30K) (*Thermo Fisher Scientific*) and purified to homogeneity on an AKTA FPLC system fitted with a 26/60 Superdex 200 pg size exclusion column (*GE Life Sciences*) equilibrated with 20 mM Tris (pH 8.0) containing 100 mM NaCl. Peak fractions as judged by the A^280^ were subjected to SDS-PAGE and the pure fractions pooled, concentrated (10 mg/ml) and stored at −80°C until further use.

The histidine tagged protein PhuS-His6 was expressed as for the non-His tagged PhuS. The lysate following removal of the cell debris was applied directly to a nickel-nitrilotriacetic acid (NTA)-agarose (*Thermo Fisher Scientific*) column (1 × 5 ml) previously equilibrated with 20 mM Tris (pH 8.0) containing 0.5 M NaCl and 5 mM imidazole. The column was washed with 10 volumes of equilibration buffer, followed by 10 volumes of wash buffer (20 mM Tris-, pH 8.0, containing 0.5 M NaCl and 60 mM imidazole), and the protein eluted in 20 mM Tris (pH 8.0) containing 0.25 M NaCl and 500 mM imidazole. The purified protein was exchanged by dialysis into 20 mM Tris (pH 8.0) containing 100 mM NaCl concentrated (10 mg/ml) and stored at −80°C until further use.

Heme solutions were prepared in 0.1 N NaOH and the pH adjusted with the identical buffer used to prepare the PhuS protein samples. Heme loading of the purified PhuS protein was carried out by addition of a 1.5:1 ratio of heme to protein. Excess heme was removed over a Sephadex G-50 column (*GE Life Sciences*) equilibrated with 20 mM Tris (pH 8.0). All buffered heme solutions were used within 20 min of preparation. Heme stock solution concentrations and the stoichiometry of the final holo-PhuS complexes were determined by pyridine hemochrome as previously described (44).

### Expression and Purification of HemO

HemO was purified as previously reported with slight modification (12). HemO lysate was applied to a Q-Sepharose Fast Flow column (2.5 cm × 6 cm) (*GE Life Sciences*) equilibrated with 20 mM Tris (pH 8.0 at 4°C), 100 mM NaCl. Protein was eluted with a 20 mM Tris (pH 8.0 at 4°C), 100−500 mM NaCl gradient. Peak protein fractions were determined SDS-PAGE and were pooled, concentrated and dialyzed against 20 mM Tris buffer (pH 8) 100 mM NaCl at 4°C. The protein (5-6 ml) was further purified by FPLC over a 26/60 Superdex 200 pg size exclusion column (*GE Life Sciences*) equilibrated with 20 mM Tris (pH 8.0) containing 100 mM NaCl. Peak fractions as judged by the A280 were subjected to SDS-PAGE and the pure fractions pooled, concentrated (10 mg/ml) and stored at −80°C until further use.

### Expression and Purification of P. aeruginosa Fur

The Fur protein was expressed and purified as previously described (45). The Fur lysate, conjugated with glutathione S-transferase (GST), was placed in a glutathione super-flow column (*Clontech*), equilibrated with 20 mM Tris-HCl (pH 8.0), and washed with 5 column volumes of the same buffer. The protein was eluted with 50 mM Tris-HCl (pH 8.0) and 33 mM glutathione, and eluted fractions were analyzed by native-PAGE. Fractions containing GST-*pa*Fur were cleaved using a Thrombin CleanCleave kit (*Sigma*-*Aldrich*). Briefly, an aliquot of thrombin-agarose resin (50% V/V) was mixed with 1 mg of GST-*pa*Fur and 100 μL of 10X cleavage buffer. The mixture was incubated at 37°C for 3h while collecting 10 μL aliquots at every hour. Fractions were measured by the A280 and checked for purity via SDS-PAGE. The fully cleaved protein was pooled and exchanged by dialysis in 20 mM Tris-HCl (pH 8.0), concentrated (10 mg/mL) and stored in −80°C until further use.

### Chromatin Immunoprecipitation (ChIP)-PCR

A single isolated colony of *Pseudomonas* PAO1 or Δ*phuS* strains was used to inoculated 10 mL of LB broth, and grown overnight at 37°C. The bacteria were then harvested and resuspended in 2 mL of M9 minimal medium. The resuspended cultures were used to inoculate 25 mL of fresh M9 low-iron medium to a starting A600 of 0.04. Cultures were grown at 37°C with shaking for 5 h. Cells were harvested at 7,000 × g for 3 min (25°C) and resuspended in 2 mL of M9 minimal media that was used to inoculate 25 mL of M9 medium to a starting A600 of 0.04. Cultures were grown for 3 hr in iron-limiting conditions before the addition of 0.5 μM heme. Following an additional 2 h, cells were harvested at 7,000 × g for 10 min, and resuspended in 2 mL of phosphate buffered saline (PBS). The aliquots were treated with formaldehyde to 1% (V/V). The cells were gently agitated at room temperature for 10 min, and then the crosslinking was quenched with glycine to a final concentration of 10 mg/mL. Cells were then gently shaken at 4°C for 30 min, centrifuged, and washed twice with PBS. Finally, cells were resuspended in 2 mL of lysis buffer (100 mM Tris (pH 8.0) containing 300 mM NaCl, 10 mM EDTA, 0.1 mM PMSF and 50 μg/mL lysozyme), mixed 10 min at 4°C and then sonicated (50 sec with 5 sec pulse 1 min pause at 80% amplitude) before its centrifugation to remove cell debris. Cell extracts were aliquoted into 1 mL volumes and frozen at −80°C until further use. Magnetic beads conjugated with IgG Protein A/G (*New England Biolabs*) were pre-blocked with 0.5 mg/mL of sonicated salmon sperm DNA (*Thermo Scientific*) and bovine serum albumin (*Sigma Aldrich*) and washed with 100 mM Tris (pH 8.0) containing 300 mM NaCl to create a slurry. Lysates were precleared with 50 μL of the bead slurry per 500 μL of cell lysate and incubated with gentle agitation at room temperature for 1 h, followed by centrifugation for 5 min at 2500 × g (4°C). The supernatant was collected, and total protein concentration was measured via the bicinchoninic (BCA) assay (*BioRad*). Samples were split into 2 × 500 μL and 2 μL anti-PhuS serum was added to one sample, both samples were then mixed by rotation at 4°C overnight. Antibodies were obtained from *Covance Custom Antibodies* and generated from purified proteins supplied by our laboratory. Antibody specificity and sensitivity was previously determined with the respective purified proteins. 100 μL washed-bead slurry was added to all samples and mixed for 30 minutes at 4°C, and the samples were centrifuged as before. Supernatant from the negative control was saved to use as input DNA. The protein-DNA complex was washed with 1X PBS several times and eluted with 0.1-0.2 M of Glycine-HCl buffer (pH 2.5-3.0). The elution was neutralized by addition of 1 M Tris buffer (pH 8). The protein-DNA complex was treated with 1 U/μl DNase I (*Novagen*) to digest non-specific DNA. The protein-DNA complex was un-crosslinked by adding 0.2 M of NaCl and incubating overnight at 65°C. The DNA remaining was purified using the QIAquick PCR Purification kit (*Qiagen*) following the manufacturer’s instructions. Specific primers designed within the *prrF1,F2* promoter region were used to amplify the isolated fragment (Table S2). PCR products were analyzed via agarose electrophoresis and visualized under UV light in an AlphaImager gel doc (*Protein Simple*). PCR amplified products were sequenced to confirm specificity.

Pull-downs with purified PhuS-His6 and genomic DNA (gDNA) of *P. aeruginosa* were performed as follows. gDNA (3 μg) was digested by *Hpa*II (*New England Biolabs*) at 37°C for 1 hr. The digested samples were loaded on an agarose gel to detect and purify fragments ranging from 100-500 bps using the Monarch^®^ DNA Gel Extraction Kit (*New England Biolabs*). The fragmented gDNA was incubated for 1 hr with 10 μM PhuS in 50 mM Tris-HCl (pH 8.0), 100 mM NaCl, 1 mM PMSF, and 1 complete mini protease inhibitor tablet (*Roche*) on a table-top roto shaker (*Scientific Industries*). 100 μl Ni-NTA resin (*Themo Fisher Scientific*) was added to the mixture and incubated for 5 min at 4°C. The resin was centrifuged at max speed (~10,000 × g) for 1 min and 3 times with 50 mM Tris-HCl (pH 8.0) containing 100 mM NaCl. PhuS-His_6_ was eluted from the resin with 50 mM Tris-HCl (pH 8.0) containing 100 mM NaCl, and 250 mM imidazole. The eluant was treated with DNase I (1 U/μl) to digest non-specific DNA and the bound DNA was purified from the complex via PCR Qiagen Kit (*Qiagen*). The purified DNA was amplified with primer sets specific to the *prrF1,F2* promoter (Table S2) analyzed via agarose electrophoresis and sequenced as described above.

### Electrophoretic Mobility Shift Assay (EMSA)

DNA fragments for EMSA experiments were obtained by annealing the 5’-biotin labeled oligonucleotides (Table S2). Sense and anti-sense oligonucleotides were annealed by mixing a 1:1 ratio, incubating at 95°C for 5 minutes followed by cooling down to room temperature. DNA was cleaned up with the QIAquick Nucleotide Removal Kit (*Qiagen*), and the concentration measured by UV absorption at 260 nm in a NanoDrop 2000c Spectrophotometer (*Thermo Scientific*). All protein oligonucleotide binding reactions were assayed in 100 mM Tris (pH 8.0) containing 500 mM NaCl, 10% glycerol and 10 ng/mL salmon sperm DNA. Apo-PhuS protein concentrations ranged from 0.1-25 μM. For the HemO titration reactions, holo-PhuS was fixed at 10 μM. HemO protein concentrations were varied across a range 0-12 molar equivalents. All reactions were incubated at 37°C for 20 min before the addition of the biotinylated probe. For all reactions, the oligonucleotides were used at a fixed concentration of 30 pM in a final volume of 10 μL. The reactions were incubated for a further 20 min at 37°C and analyzed on an 8% Tris-Glycine acrylamide native gel. The gel was pre-run in 1X Tris-Glycine buffer (pH 8.3) for 1h at 200 V 4°C and run 2h at same voltage. DNA was transferred to positively charged nylon membrane (BrightStar-Plus Positively Charged Nylon Membranes; *Invitrogen*) using a Semidry Electroblotting System (*Thermo Scientific*) with 1X Tris-Glycine (pH 8.3) for 30 min at 300 mA. Membranes were washed in 2X saline-sodium citrate buffer (SSC) (*Thermo Scientific*), for 5 min at room temperature, and DNA immobilized by UV crosslinking. The position of the Nucleic acids was visualized by chemiluminescent detection using the Chemiluminescent Nucleic Acid Detection Module (*Thermo Scientific*) following manufacturer’s instructions and exposed to X-ray film (Amersham hyperfilm ECL; *Amersham*).

### Fluorescence Anisotropy (FA)

Binding of Fur or apo-PhuS to the 5’fluorescein (FAM)-oligonucleotides (Table S1) was assessed using fluorescence anisotropy as previously described (20). Briefly, 5’-FAM oligonucleotides designed within the *prrF1/prrF2* promoter region (Table S1) were analyzed by UV-vis spectroscopy to quantify the percentage of fluorescein tag (Table 1). Double-stranded oligonucleotides were obtained by combining a 1:1 ratio of the 5’-labeled sense and antisense oligonucleotides in deionized H2O. To facilitate annealing, mixtures were heated to 95°C, 5 min, and cooled down to room temperature. The labeled double stranded oligonucleotides were stored at −80°C until further use. In a quartz cuvette, 10 nM of 5’-FAM oligonucleotide was diluted in 20 mM Tris-HCl (pH 8.0), containing 100 mM NaCl, and 0.05 mg/mL bovine serum albumin in a final volume of 500 ul. For runs with Fur, 10 nM of 5’-FAM oligonucleotide was diluted in 10 mM Bis-Tris-borate (pH 7.5), 40 mM KCl, 0.1 mM MnSO_4_, 0.1 mg/mL BSA and 10% glycerol. All measurements were performed on a K2 spectrofluorometer (*ISS*) configured in the L-format, with excitation/emission wavelengths and band widths of 495 nm and 2nm and 519 nm and 1nm, respectively. A measurement of the maximum anisotropy was performed on the 5’-FAM-oligonucleotide, the change in anisotropy was measured as a function of increasing concentrations of apo-PhuS or Fur. The addition of protein continued until no further change in anisotropy was observed. The data was fit by converting the anisotropy, *r*, to fraction bound, *F*_bound_ (the fraction of protein bound to the oligonucleotide at a given DNA concentration), using the following equation:

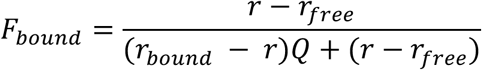

where *r*_free_ is the anisotropy of the fluorescein-labeled oligonucleotide and *r*_bound_ is the anisotropy of the oligonucleotide-protein complex at saturation. The quantum yield designated as *Q* is calculated from the changes in fluorescence intensity that occurs over the course of the experiment (*I*_bound_/ *I*_*free*_). *F*_bound_ was then plotted against the protein concentration using a one-site binding model:

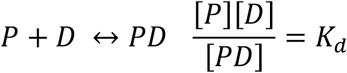

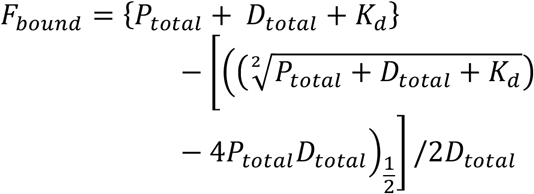

where *P* is the protein concentration and *D* is the DNA concentration. All concentrations and fluorescence changes were done in triplicate and corrected for volume changes.

For the apo-HemO titrations the experiments were performed as described above with a fixed concentration of 5’-FAM labeled *prrF*1-50 (10 nM) and holo-PhuS (1 μM). The change in anisotropy was recorded on addition of increasing concentrations of apo-HemO and converted to fraction bound and plotted against HemO molar equivalents.

## FOOTNOTES

This work was supported by National Institutes of Health Grant R01 AI134886 (to A. W.) TW wishes to acknowledge previous support from a Chemistry/Biology Interface Training Program (NIGMS/NIH T32GM066706). AW would like to thank Amanda Oglesby for the kind gift of the pGST-*paFur* expression vector.

## ABBREVIATIONS

PAO1: *Pseudomonas aeruginosa* O1
PrrF: *Pseudomonas*
RNA: responsive to iron
PrrH: *Pseudomonas*
RNA: responsive to heme
Has: Heme assimilation system
Phu: *Pseudomonas* heme uptake
BVIX: biliverdin
ECF: extra cytoplasmic function
ABC-transport system: ATP-dependent binding cassette transport system
FA: Fluorescence anisotropy

## AUTHOR CONTRIBUTIONS

AW and TW conceived and designed the study. TW performed all experiments including EMSA and FA experiments, ChIP-PCR and qPCR. SM constructed the *phuSH212R* allelic strain.TW, SM and AW interpreted and analyzed the data. TW and AW wrote the manuscript.

## REFERENCES

1. Johnson, E. E., and Wessling-Resnick, M. (2012) Iron metabolism and the innate immune response to infection. Microbes Infect 14, 207–216

2. Hood, M. I., and Skaar, E. P. (2012) Nutritional immunity: transition metals at the pathogen-host interface. Nat Rev Microbiol 10, 525–537

3. Letoffe, S., Delepelaire, P., and Wandersman, C. (2004) Free and hemophore-bound heme acquisitions through the outer membrane receptor HasR have different requirements for the TonB-ExbB-ExbD complex. J Bacteriol 186, 4067–4074

4. Contreras, H., Chim, N., Credali, A., and Goulding, C. W. (2014) Heme uptake in bacterial pathogens. Curr Opin Chem Biol 19, 34–41

5. Huang, W., and Wilks, A. (2017) Extracellular Heme Uptake and the Challenge of Bacterial Cell Membranes. Annu Rev Biochem 86, 799–823

6. Ochsner, U. A., Johnson, Z., and Vasil, M. L. (2000) Genetics and regulation of two distinct haem-uptake systems, *phu* and *has*, in *Pseudomonas aeruginosa*. Microbiology 146 (Pt 1), 185–198

7. Smith, A. D., and Wilks, A. (2015) Differential contributions of the outer membrane receptors PhuR and HasR to heme acquisition in *Pseudomonas aeruginosa*. J Biol Chem 290, 7756–7766

8. Mascher, T. (2013) Signaling diversity and evolution of extracytoplasmic function (ECF) sigma factors. Curr Opin Microbiol 16, 148–155

9. Helmann, J. D. (2002) The extracytoplasmic function (ECF) sigma factors. Adv Microb Physiol 46, 47–110

10. Lansky, I. B., Lukat-Rodgers, G. S., Block, D., Rodgers, K. R., Ratliff, M., and Wilks, A. (2006) The cytoplasmic heme-binding protein (PhuS) from the heme uptake system of *Pseudomonas aeruginosa* is an intracellular heme-trafficking protein to the delta-regioselective heme oxygenase. J Biol Chem 281, 13652–13662

11. O’Neill, M. J., and Wilks, A. (2013) The *P. aeruginosa* heme binding protein PhuS is a heme oxygenase titratable regulator of heme uptake. ACS Chem Biol 8, 1794–1802

12. Ratliff, M., Zhu, W., Deshmukh, R., Wilks, A., and Stojiljkovic, I. (2001) Homologues of neisserial heme oxygenase in gram-negative bacteria: degradation of heme by the product of the *pigA* gene of *Pseudomonas aeruginosa*. J Bacteriol 183, 6394–6403

13. Dent, A. T., Mourino, S., Huang, W., and Wilks, A. (2019) Post-transcriptional regulation of the *Pseudomonas aeruginosa* heme assimilation system (Has) fine-tunes extracellular heme sensing. J Biol Chem 294, 2771–2785

14. Wilderman, P. J., Sowa, N. A., FitzGerald, D. J., FitzGerald, P. C., Gottesman, S., Ochsner, U. A., and Vasil, M. L. (2004) Identification of tandem duplicate regulatory small RNAs in *Pseudomonas aeruginosa* involved in iron homeostasis. Proc Natl Acad Sci U S A 101, 9792–9797

15. Oglesby, A. G., Farrow, J. M., 3rd, Lee, J. H., Tomaras, A. P., Greenberg, E. P., Pesci, E. C., and Vasil, M. L. (2008) The influence of iron on *Pseudomonas aeruginosa* physiology: a regulatory link between iron and quorum sensing. J Biol Chem 283, 15558–15567

16. Reinhart, A. A., Powell, D. A., Nguyen, A. T., O’Neill, M., Djapgne, L., Wilks, A., Ernst, R. K., and Oglesby-Sherrouse, A. G. (2015) The *prrF*-encoded small regulatory RNAs are required for iron homeostasis and virulence of *Pseudomonas aeruginosa*. Infect Immun 83, 863–875

17. Oglesby-Sherrouse, A. G., and Vasil, M. L. (2010) Characterization of a heme-regulated non-coding RNA encoded by the *prrF* locus of *Pseudomonas aeruginosa*. PLoS One 5, e9930

18. Reinhart, A. A., Nguyen, A. T., Brewer, L. K., Bevere, J., Jones, J. W., Kane, M. A., Damron, F. H., Barbier, M., and Oglesby-Sherrouse, A. G. (2017) The *Pseudomonas aeruginosa* PrrF Small RNAs Regulate Iron Homeostasis during Acute Murine Lung Infection. Infect Immun 85 e00764–16

19. Kaur, A. P., Lansky, I. B., and Wilks, A. (2009) The role of the cytoplasmic heme-binding protein (PhuS) of *Pseudomonas aeruginosa* in intracellular heme trafficking and iron homeostasis. J Biol Chem 284, 56–66

20. Kaur, A. P., and Wilks, A. (2007) Heme inhibits the DNA binding properties of the cytoplasmic heme binding protein of *Shigella dysenteriae* (ShuS). Biochemistry 46, 2994–3000

21. Deredge, D. J., Huang, W., Hui, C., Matsumura, H., Yue, Z., Moenne-Loccoz, P., Shen, J., Wintrode, P. L., and Wilks, A. (2017) Ligand-induced allostery in the interaction of the *Pseudomonas aeruginosa* heme binding protein with heme oxygenase. Proc Natl Acad Sci U S A 114, 3421–3426

22. Mourino, S., Giardina, B. J., Reyes-Caballero, H., and Wilks, A. (2016) Metabolite-driven Regulation of Heme Uptake by the Biliverdin IXbeta/delta-Selective Heme Oxygenase (HemO) of *Pseudomonas aeruginosa*. J Biol Chem 291, 20503–20515

23. Dent, A. T., and Wilks, A. (2020) Contributions of the heme coordinating ligands of the *Pseudomonas aeruginosa* outer membrane receptor HasR to extracellular heme sensing and transport. J Biol Chem 295, 10456–10467

24. Mey, A. R., Craig, S. A., and Payne, S. M. (2005) Characterization of *Vibrio cholerae* RyhB: the RyhB regulon and role of *ryhB* in biofilm formation. Infect Immun 73, 5706–5719

25. Murphy, E. R., and Payne, S. M. (2007) RyhB, an iron-responsive small RNA molecule, regulates *Shigella dysenteriae* virulence. Infect Immun 75, 3470–3477

26. Jacques, J. F., Jang, S., Prevost, K., Desnoyers, G., Desmarais, M., Imlay, J., and Masse, E. (2006) RyhB small RNA modulates the free intracellular iron pool and is essential for normal growth during iron limitation in *Escherichia coli*. Mol Microbiol 62, 1181–1190

27. Leclerc, J. M., Dozois, C. M., and Daigle, F. (2013) Role of the *Salmonella enterica* serovar Typhi Fur regulator and small RNAs RfrA and RfrB in iron homeostasis and interaction with host cells. Microbiology (Reading) 159, 591–602

28. Nguyen, A. T., O’Neill, M. J., Watts, A. M., Robson, C. L., Lamont, I. L., Wilks, A., and Oglesby-Sherrouse, A. G. (2014) Adaptation of iron homeostasis pathways by a *Pseudomonas aeruginosa* pyoverdine mutant in the cystic fibrosis lung. J Bacteriol 196, 2265–2276

29. Marvig, R. L., Damkiaer, S., Khademi, S. M., Markussen, T. M., Molin, S., and Jelsbak, L. (2014) Within-host evolution of *Pseudomonas aeruginosa* reveals adaptation toward iron acquisition from hemoglobin. mBio 5, e00966–00914

30. Cournac, A., and Plumbridge, J. (2013) DNA looping in prokaryotes: experimental and theoretical approaches. J Bacteriol 195, 1109–1119

31. Roncarati, D., Pelliciari, S., Doniselli, N., Maggi, S., Vannini, A., Valzania, L., Mazzei, L., Zambelli, B., Rivetti, C., and Danielli, A. (2016) Metal-responsive promoter DNA compaction by the ferric uptake regulator. Nat Commun 7, 1–13

32. Agriesti, F., Roncarati, D., Musiani, F., Del Campo, C., Iurlaro, M., Sparla, F., Ciurli, S., Danielli, A., and Scarlato, V. (2014) FeON-FeOFF: the *Helicobacter pylori* Fur regulator commutates iron-responsive transcription by discriminative readout of opposed DNA grooves. Nucleic Acids Res 42, 3138–3151

33. Carpenter, B. M., Gancz, H., Benoit, S. L., Evans, S., Olsen, C. H., Michel, S. L., Maier, R. J., and Merrell, D. S. (2010) Mutagenesis of conserved amino acids of *Helicobacter pylori* fur reveals residues important for function. J Bacteriol 192, 5037–5052

34. Little, A. S., Okkotsu, Y., Reinhart, A. A., Damron, F. H., Barbier, M., Barrett, B., Oglesby-Sherrouse, A. G., Goldberg, J. B., Cody, W. L., Schurr, M. J., Vasil, M. L., and Schurr, M. J. (2018) *Pseudomonas aeruginosa* AlgR Phosphorylation Status Differentially Regulates Pyocyanin and Pyoverdine Production. mBio 9,699–718

35. Cody, W. L., Pritchett, C. L., Jones, A. K., Carterson, A. J., Jackson, D., Frisk, A., Wolfgang, M. C., and Schurr, M. J. (2009) *Pseudomonas aeruginosa* AlgR controls cyanide production in an AlgZ-dependent manner. J Bacteriol 191, 2993–3002

36. Morici, L. A., Carterson, A. J., Wagner, V. E., Frisk, A., Schurr, J. R., Honer zu Bentrup, K., Hassett, D. J., Iglewski, B. H., Sauer, K., and Schurr, M. J. (2007) *Pseudomonas aeruginosa* AlgR represses the Rhl quorum-sensing system in a biofilm-specific manner. J Bacteriol 189, 7752–7764

37. Okkotsu, Y., Tieku, P., Fitzsimmons, L. F., Churchill, M. E., and Schurr, M. J. (2013) *Pseudomonas aeruginosa* AlgR phosphorylation modulates rhamnolipid production and motility. J Bacteriol 195, 5499–5515

38. Whitchurch, C. B., Erova, T. E., Emery, J. A., Sargent, J. L., Harris, J. M., Semmler, A. B., Young, M. D., Mattick, J. S., and Wozniak, D. J. (2002) Phosphorylation of the *Pseudomonas aeruginosa* response regulator AlgR is essential for type IV fimbria-mediated twitching motility. J Bacteriol 184, 4544–4554

39. Kong, W., Zhao, J., Kang, H., Zhu, M., Zhou, T., Deng, X., and Liang, H. (2015) ChIP-seq reveals the global regulator AlgR mediating cyclic di-GMP synthesis in *Pseudomonas aeruginosa*. Nucleic Acids Res 43, 8268–8282

40. Mohr, C. D., Sonsteby, S. K., and Deretic, V. (1994) The *Pseudomonas aeruginosa* homologs of *hemC* and *hemD* are linked to the gene encoding the regulator of mucoidy AlgR. Mol Gen Genet 242, 177–184

41. Hoang, T. T., Karkhoff-Schweizer, R. R., Kutchma, A. J., and Schweizer, H. P. (1998) A broad-host-range Flp-FRT recombination system for site-specific excision of chromosomally-located DNA sequences: application for isolation of unmarked *Pseudomonas aeruginosa* mutants. Gene 212, 77–86

42. Barker, K. D., Barkovits, K., and Wilks, A. (2012) Metabolic flux of extracellular heme uptake in *Pseudomonas aeruginosa* is driven by the iron-regulated heme oxygenase (HemO). J Biol Chem 287, 18342–18350

43. O’Neill, M. J., Bhakta, M. N., Fleming, K. G., and Wilks, A. (2012) Induced fit on heme binding to the *Pseudomonas aeruginosa* cytoplasmic protein (PhuS) drives interaction with heme oxygenase (HemO). Proc Natl Acad Sci U S A 109, 5639–5644

44. Fuhrop, J. H., and Smith, K. M. (eds). (1975) Porphyrins and Metalloporphyrins, pp 804–807, Elsevier, Amsterdam

45. Pohl, E., Haller, J. C., Mijovilovich, A., Meyer-Klaucke, W., Garman, E., and Vasil, M. L. (2003) Architecture of a protein central to iron homeostasis: crystal structure and spectroscopic analysis of the ferric uptake regulator. Mol Microbiol 47, 903–915

